# SC3 - consensus clustering of single-cell RNA-Seq data

**DOI:** 10.1101/036558

**Authors:** Vladimir Yu. Kiselev, Kristina Kirschner, Michael T. Schaub, Tallulah Andrews, Andrew Yiu, Tamir Chandra, Kedar N Natarajan, Wolf Reik, Mauricio Barahona, Anthony R Green, Martin Hemberg

**Affiliations:** Wellcome Trust Sanger Institute, Hinxton, Cambridge, UK; Cambridge Institute for Medical Research, Wellcome Trust/MRC Stem Cell Institute and Department of Haematology, University of Cambridge, Hills Road, Cambridge, UK; Department of Mathematics and naXys, University of Namur, Belgium; ICTEAM, Université catholique de Louvain, Belgium; Epigenetics Programme, The Babraham Institute, Babraham, Cambridge, UK; EMBL-European Bioinformatics Institute, Hinxton, Cambridge, UK; Centre for Trophoblast Research, University of Cambridge, Cambridge, UK; Department of Mathematics, Imperial College London, London, UK

## Abstract

Using single-cell RNA-seq (scRNA-seq), the full transcriptome of individual cells can be acquired, enabling a quantitative cell-type characterisation based on expression profiles. However, due to the large variability in gene expression, identifying cell types based on the transcriptome remains challenging. We present Single-Cell Consensus Clustering (SC3), a tool for unsupervised clustering of scRNA-seq data. SC3 achieves high accuracy and robustness by consistently integrating different clustering solutions through a consensus approach. Tests on twelve published datasets show that SC3 outperforms five existing methods while remaining scalable, as shown by the analysis of a large dataset containing 44,808 cells. Moreover, an interactive graphical implementation makes SC3 accessible to a wide audience of users, and SC3 aids biological interpretation by identifying marker genes, differentially expressed genes and outlier cells. We illustrate the capabilities of SC3 by characterising newly obtained transcriptomes from subclones of neoplastic cells collected from patients.

## Introduction

With the recent advent of single cell RNA-seq (scRNA-seq) technology, researchers are now able to quantify the entire transcriptome of individual cells, opening up a wide spectrum of biological applications. One key application of scRNA-seq is the ability to determine cell types based on their transcriptome profile alone^1–3^. The diversity of cell-types is a fundamental property of higher eukaryotes. Traditionally, cell type was defined based on shape and morphological properties, but tools from molecular biology have enabled researchers to categorise cells based on surface markers^4,5^. However, morphology or a small number of marker proteins are not sufficient to characterise complex cellular phenotypes. scRNA-seq opens up the possibility to group cells based on their genome-wide transcriptome profiles, which is likely to provide a better representation of the cellular phenotype. Indeed, several studies have already used scRNA-seq to identify novel cell-types^1–3^, demonstrating its potential to unravel and catalogue the full diversity of cells in the human body.

A full characterisation of the transcriptional landscape of individual cells holds an enormous potential, both for basic biology and clinical applications. An important medical application is to study cancer, widely known to be a heterogeneous disease with multiple subclones coexisting within the same tumor. Until recently, tumor heterogeneity has mainly been assessed at the DNA level by genome sequencing^6–8^. The use of scRNA-seq makes it possible to characterise the transcriptional landscape of different subclones within the same tumour. A better understanding of the transcriptome in the different subclones could thus yield important insights about drug resistance, thereby informing the development of novel therapies. However, due to the large variability in gene expression, identifying subclones from patient transcriptomes remains challenging^9,10^

Mathematically, the problem of *de novo* identification of cell-types from data may be seen as an unsupervised clustering problem, i.e., how to separate cells into non-overlapping groups without a *priori* information as to the number of groups and group labels. However, the lack of training data and reliable benchmarks for validation renders this unsupervised clustering a hard problem. For scRNA-seq the challenge is further compounded by technical errors and biases that remain incompletely understood, a high degree of *biological* variability in gene expression^11^, and the high dimensionality of the transcriptome.

Although scRNA-seq is a relatively new technology, several groups have developed custom clustering methods for single cell data^2,12–16^. Yet, these clustering methods have one or more of the following shortcomings: (i) they have not been thoroughly benchmarked against each other and standard algorithms; (ii) it is not clear how they can be scaled to large datasets; (iii) there is no interactive, user-friendly implementation that includes support to facilitate the biological interpretation of the clusters; and (iv) the number of clusters, *k*, has to be fixed a *priori* by the user and there is no support to explore different hierarchies of clusters. The last point is particularly relevant when studying complex biological tissues as previous studies have found biologically meaningful cell populations at several levels of granularity^14,17,18^

We present a novel interactive clustering tool for scRNA-seq data, SC3 (**S**ingle **C**ell **C**onsensus **C**lustering) a user-friendly R-package with a graphical interface. The main innovations of SC3 is the demonstration that *accurate and robust* results can be obtained by combining several well-established techniques using a consensus clustering approach. The latter aspect is particularly important in this context, as it facilitates *reproducible* analyses from inherently noisy scRNAseq data. In addition, SC3 has several features to facilitate the evaluation of the clustering quality, and to aid the user in determining the appropriate number of clusters. We demonstrate the performance of SC3 by applying it to twelve published datasets and thoroughly benchmarking the method against five other methods. Furthermore, we showcase the scalability of SC3 to very large datasets by analysing a dataset with ~45,000 cells^15^. Crucially, in addition to providing cell clusters, SC3 includes several features for integration into bioinformatics analysis pipelines and for facilitating biological interpretation. These features include tSNE plots, identifying marker genes and differentially expressed genes, detection of outlier cells, and an integrated link to Gene Ontology analysis.

We apply SC3 to the first single cell RNA-Seq data from haematopoietic stem cells isolated directly from patients with myeloproliferative neoplasm. Myeloproliferative neoplasms are a heterogeneous disease, with each patient typically harbouring multiple neoplastic subclones that coexist for long periods of time^19^. Therefore, transcriptome data from patients are hard to interpret due to its inherent heterogeneity within these cells. Here, we identify and validate clusters corresponding to different subclones within two patients with different mutational landscapes. We also characterise the differences between their expression profiles.

## Results

### Consensus clustering as a robust methodology

The output of a scRNA-seq experiment is typically represented as an expression matrix consisting of *g* rows (corresponding to genes or transcripts) and *N* columns (corresponding to cells). SC3 takes this expression matrix as its input. Note that SC3 does not carry out quality control, normalisation or batch correction internally, and it is important that the user has filtered the input data appropriately to ensure that the clustering is not confounded by technical artifacts. The SC3 algorithm consists of five steps: (i) a gene filter, (ii) distance calculations, (iii) transformations combined with (iv)*k*-means clustering, followed by (v) the consensus step (Fig. 1a, Methods). Note, in particular, that the distance calculations reflect a change of coordinate space, as we go from the expression matrix (*g* × *N*) to a cell-to-cell matrix (*N* × *N*). In the transformation step we modify this cell by cell matrix by keeping only the first *d* principal components of the distance matrix (or eigenvectors of the associated graph Laplacian) for the final clustering. Each of the above steps requires, in principle, the specification of a number of parameters, e.g. different metrics to calculate distances between the cells, or the particular type of transformation. Choosing such optimal parameter values is difficult and time-consuming, in general. To avoid this problem, SC3 utilizes a parallelisation approach, whereby a significant subset of the parameter space is evaluated simultaneously to obtain a set of clusterings. Instead of trying to pick the optimal solution from this set, we combine *all* the different clustering outcomes into a consensus matrix that summarises how often each pair of cells is located in the same cluster. The final result provided by SC3 is determined by complete-linkage hierarchical clustering of the consensus matrix into *k* groups. Using this approach, we can leverage the power of a plentitude of well-established methods, while additionally gaining robustness against the particularities of any single method.

**Figure 1.**
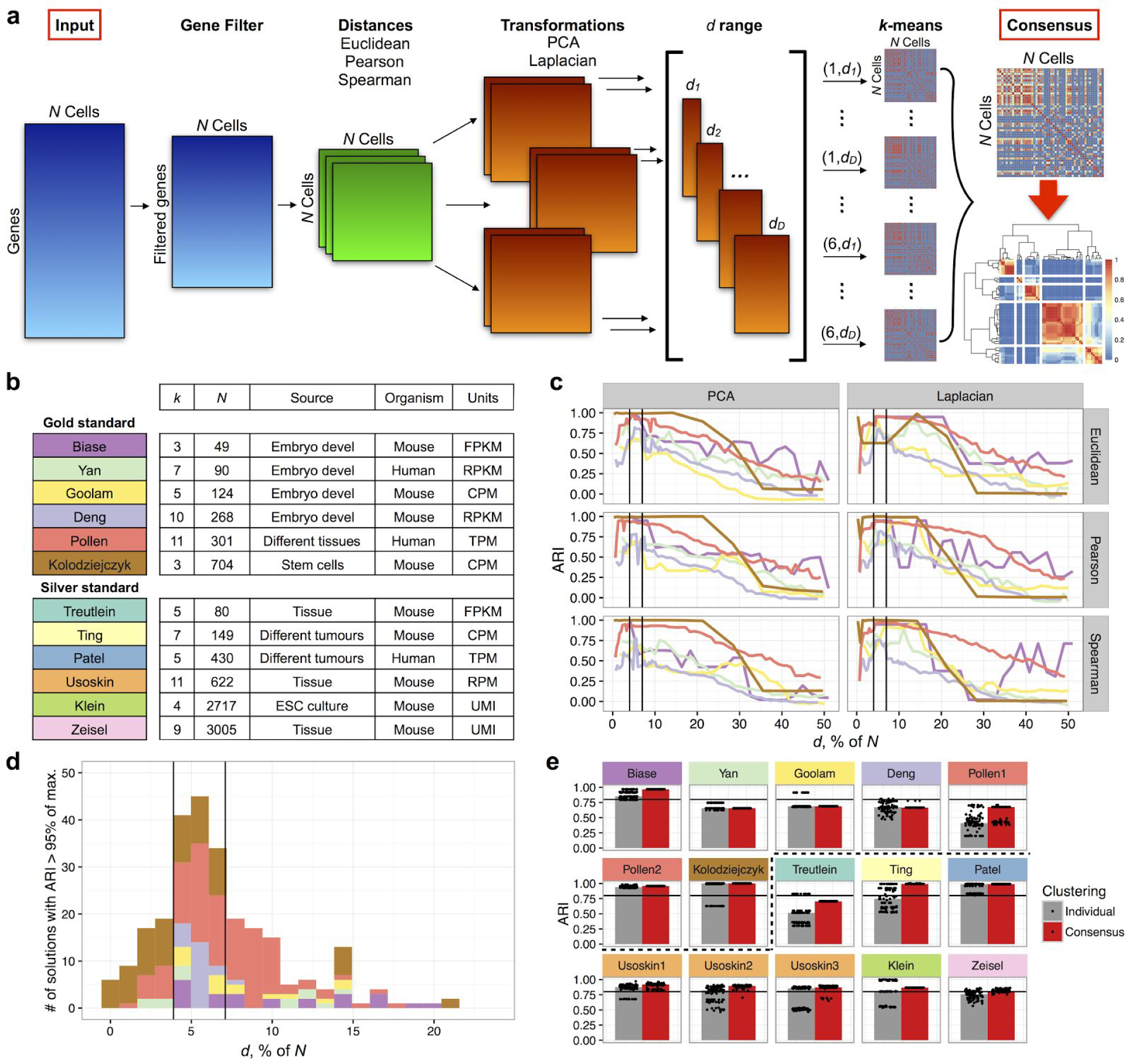
The SC3 framework for consensus clustering. **(a)** Overview of clustering with SC3 framework (see Methods). A total of 6*D* clusterings are obtained, where *D* is the total number of dimensions *d*_1_, …, *d_D_* considered. These clusterings are then combined through a consensus step to increase accuracy and robustness. Here, the consensus step is exemplified using the Treutlein data: the binary matrices (Methods) corresponding to each clustering are averaged, and the resulting matrix is segmented using hierarchical clustering up to the *k*th hierarchical level (*k* = 5 in this example). (**b**) Published datasets used to set SC3 parameters. *N* is the number of cells in a dataset; *k* is the number of clusters originally identified by the authors^9,14,17,18,20–23,63–66^; Units: RPKM is Reads Per Kilobase of transcript per Million mapped reads, RPM is Reads Per Million mapped reads, FPKM is Fragments Per Kilobase of transcript per Million mapped reads, TPM is Transcripts Per Million mapped reads. (**c**) Testing the distances, nonlinear transformations and *d* range. Median of ARI over 100 realizations of the SC3 clustering for six gold standard datasets (Biase, Yan, Goolam, Kolodziejczyk, Deng and Pollen, colours as in (**b**)). The x-axis shows the number of eigenvectors *d* (see (**a**)) as a percentage of the total number of cells, *N*. The black vertical lines indicate the interval *d* = 4-7% of the total number of cells *N*, showing high accuracy in the classification, **(d)** Histogram of the *d* values where ARI>.95 is achieved for the gold standard datasets. The black vertical lines indicate the same as in (**c**). (**e**) 100 realizations of the SC3 clustering of the datasets shown in (**b**). *Individual* corresponds to clustering without consensus step. *Consensus* corresponds to the consensus clustering over the parameter set (Methods). The black line corresponds to ARI=0.8. Dots represent individual clustering runs. The dashed black line separates gold and silver standard datasets.

The SC3 pipeline contains several parameters, whose ranges can be flexibly adjusted by the user in a simple manner. Note that in principle, additional clustering approaches could be included in the pipeline to be considered within the consensus step. To constrain the parameter values for our analysis here, we first considered six publicly available scRNA-Seq datasets^17,20–24^ (Fig. 1b). The datasets were selected on the basis that one can be highly confident in the cell-labels as they represent cells from different developmental stages (Biase, Deng, Yan and Goolam), stem cells grown in different conditions (Kolodziejczyk) or cell lines (Pollen), and thus we consider them as ‘gold standard’. According to the authors, the Pollen dataset contains two distinct hierarchies and the cells can be grouped either into 4 or 11 clusters.

To quantify the similarity between the reference labels and the clusters obtained by SC3, we used the Adjusted Rand Index (ARI, see Methods) which ranges from 1, when the clusterings are identical, to 0 when the similarity is what one would expect by chance. For the gold standard datasets, we found that clustering performance was largely unaffected by different choices of gene filter, distance metrics and transformations (Fig. 1c, S1-S7). Additionally, we investigated the impact of dropouts by considering a modified distance metric that ignores dropouts, but we found that this did not improve the performance (Fig. S9, Methods). In contrast, we found that the quality of the outcome as measured by the ARI was sensitive to the number of eigenvectors, *d*, retained after the spectral transformation: the ARI is low for small values of *d*, and then increases to its maximum before dropping close to 0 as *d* approaches *N*/2. For each of the gold standard datasets we identified the values of *d* where an ARI >95% of the maximum was obtained. For all six datasets we find that the best clusterings were typically achieved when *d* is between 4-7% of the number of cells, *N* (Fig. 1d, Methods). The robustness of the 4-7% region was supported by a simulation experiment where the six gold standard datasets were downsampled by a factor of ten (Methods and Fig. S10). We further tested the SC3 pipeline on six other published datasets, where the cell labels can only be considered ‘silver standard’ since they were assigned using computational methods and the authors’ knowledge of the underlying biology. Again, we find that SC3 performs well when using *d* in the 4-7% of *N* interval (Fig. S8). The silver standard results demonstrate the SC3 is consistent with the authors’ methods, but the results could be affected by the fact that the labels were assigned using a procedure that also relied on a similar dimensionality reduction. The final step, consensus clustering improves the stability of the solution, *k*-means based methods will typically provide different outcomes depending on the initial conditions. We find that this variability is significantly reduced with the consensus approach. In some cases we even find that the consensus solution is better than any of the individual solutions (Fig. 1e).

To benchmark SC3, we considered five other methods: tSNE^25^ followed by *k*-means clustering (a method similar to the one used by Grün et al^1^), pcaReduce^13^, SNN-Cliq^12^, SINCERA^16^ and SEURAT^15^ As Fig. 2a shows, SC3 performs better than the five tested methods across all datasets (Wilcoxon signed-rank test p-value < 0.01), with the exception of the Pollen1, Biase (where ARIs of other methods are similar to SC3 but slightly higher) and Yan dataset (where pcaReduce performed better). In addition to considering accuracy, we also compared the stability of SC3 with other stochastic methods (pcaReduce and tSNE+kmeans, but not SEURAT) by running them 100 times. The different outcomes are shown as black dots in Fig. 2a and in contrast to the other methods that rely on different initializations, SC3 is highly stable. Usually a single solution is obtained as indicated by our stability measure (Methods) in Fig. 2b.

**Figure 2.**
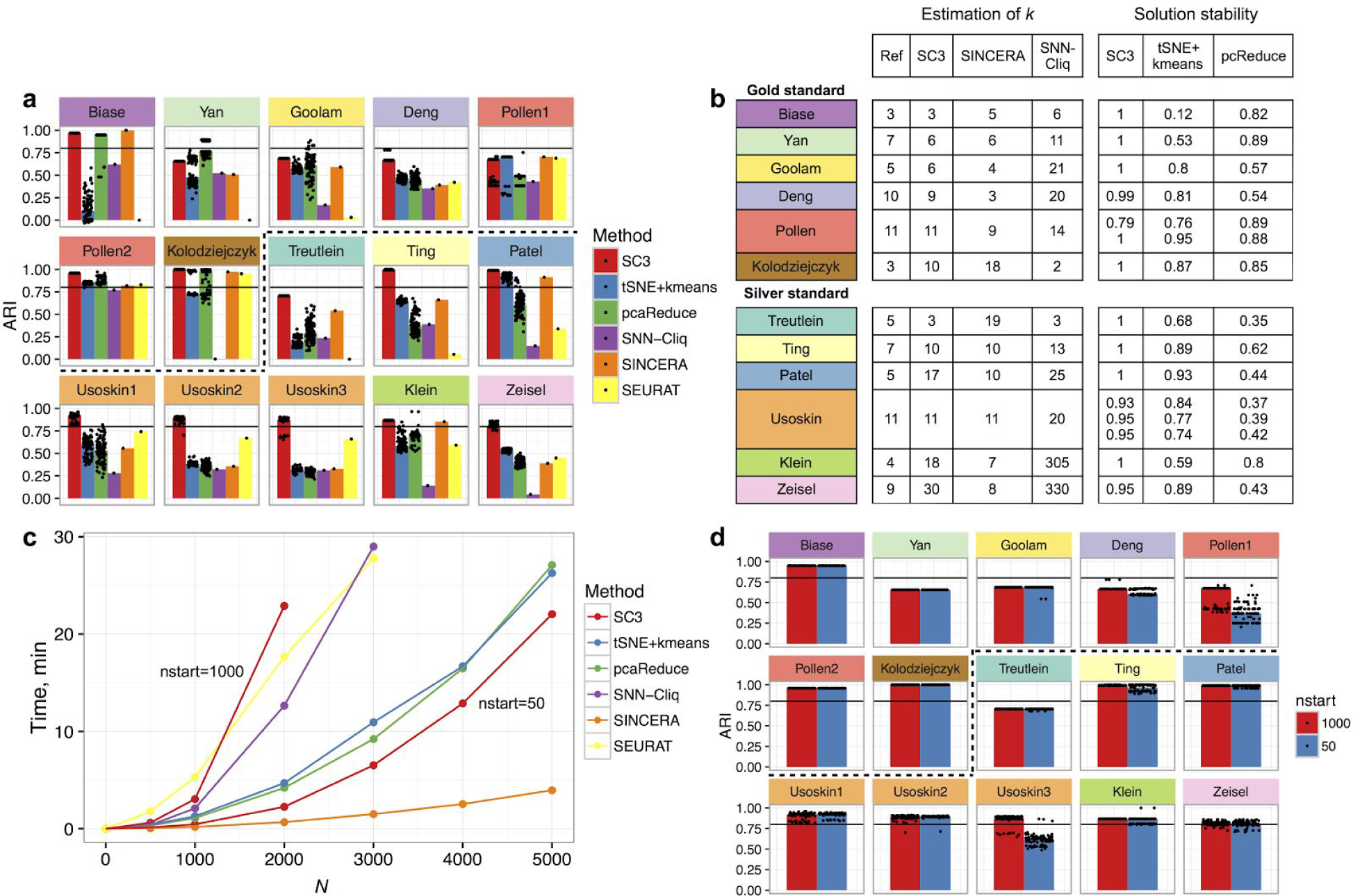
Benchmarking of SC3 against existing methods. **(a)** SC3, tSNE+kmeans and pcaReduce were applied 100 times to each dataset to evaluate accuracy and stability. SNN-Cliq and SINCERA are deterministic and were thus run only once. SEURAT was also run once, however was optimised over different values of the density parameter *G* (Methods). Each panel shows the similarity between the inferred clusterings and the reference labels. The similarity is quantified by the Adjusted Rand Index (ARI, see Methods) which ranges from 1, when the clusterings are identical, to 0 when the similarity is what one would expect by chance. The ARI was calculated for each run of the respective method (black dots). The top of each bar corresponds to the median of the distribution of the black dots. For the Pollen and Usoskin datasets we considered all the different hierarchies reported in the original papers (Pollen1 *k* = 4, Pollen2 *k* = 11, Usoskin 1 *k* = 4, Usoskin2 *k* = 8, Usoskin3 *k* = 11). The black line indicates ARI = 0.8. The dashed black line separates gold and silver standard datasets, **(b)** Number of clusters 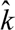 predicted by SC3, SINCERA and SNN-Cliq for all datasets. Ref is the reference clustering reported by the authors, **(c)** Run times for different clustering methods as a function of the number of cells (*N*). All methods were run on a MacBook Pro (Mid 2014), OS X Yosemite 10.10.5 with 2.8 GHz Intel Core i7 processor, 16 GB 1600 MHz DDR3 of RAM. Two results shown for SC3 correspond to nstart=1000 and nstart=50, where nstart is the number of starting points for k-means clustering, **(d)** Reducing the number of k-means runs (nstart) from 1,000 to 50 results only in a slightly worse performance for SC3, yet with significant computational savings, as shown in **(c).** The black line indicates ARI = 0.8. The dashed black line separates gold and silver standard datasets.

### SC3 can be scaled to large datasets

Although SC3’s consensus strategy provides a high accuracy, it comes at a moderate computational cost: the run time for *N*=2,000 is ˜20 mins (Fig. 2c). The main bottleneck is the k-means clustering and by reducing how many different runs are considered it is possible to cluster 5,000 cells in ~20 mins with only a slight reduction in accuracy (Fig. 2d). Unfortunately, the scaling of the run time with *N* makes it impractical to use the version of SC3 described above for datasets with >5,000 cells. To apply SC3 to larger datasets, we have implemented a hybrid approach that combines unsupervised and supervised methodologies. When the number of cells is large (*N*>5,000), SC3 selects a subset of 5,000 cells uniformly at random, and obtains clusters from this subset as described above. Subsequently, the inferred labels are used to train a support vector machine (SVM, Methods), which is employed to assign labels to the remaining cells. Training the SVM typically takes only a few minutes, thus allowing SC3 to be applied to very large datasets.

To test the SVM based approach in isolation, we used the gold standard datasets from Fig. 1c with the authors’ reference labels to train the SVM. Our results demonstrate that using <20% of the cells for training it is possible to accurately predict the labels of the remaining cells (Fig. S11). For the situation where labels are assigned using the unsupervised clustering described above, we find that it is still possible to achieve an ARI>0.8 with only 20% of the training cells for five out of seven datasets (Fig. 3a). Taken together, the result shows that the use of an SVM to predict cell labels works well - the loss of accuracy is mainly due to the use of fewer cells in the training set.

**Figure 3.**
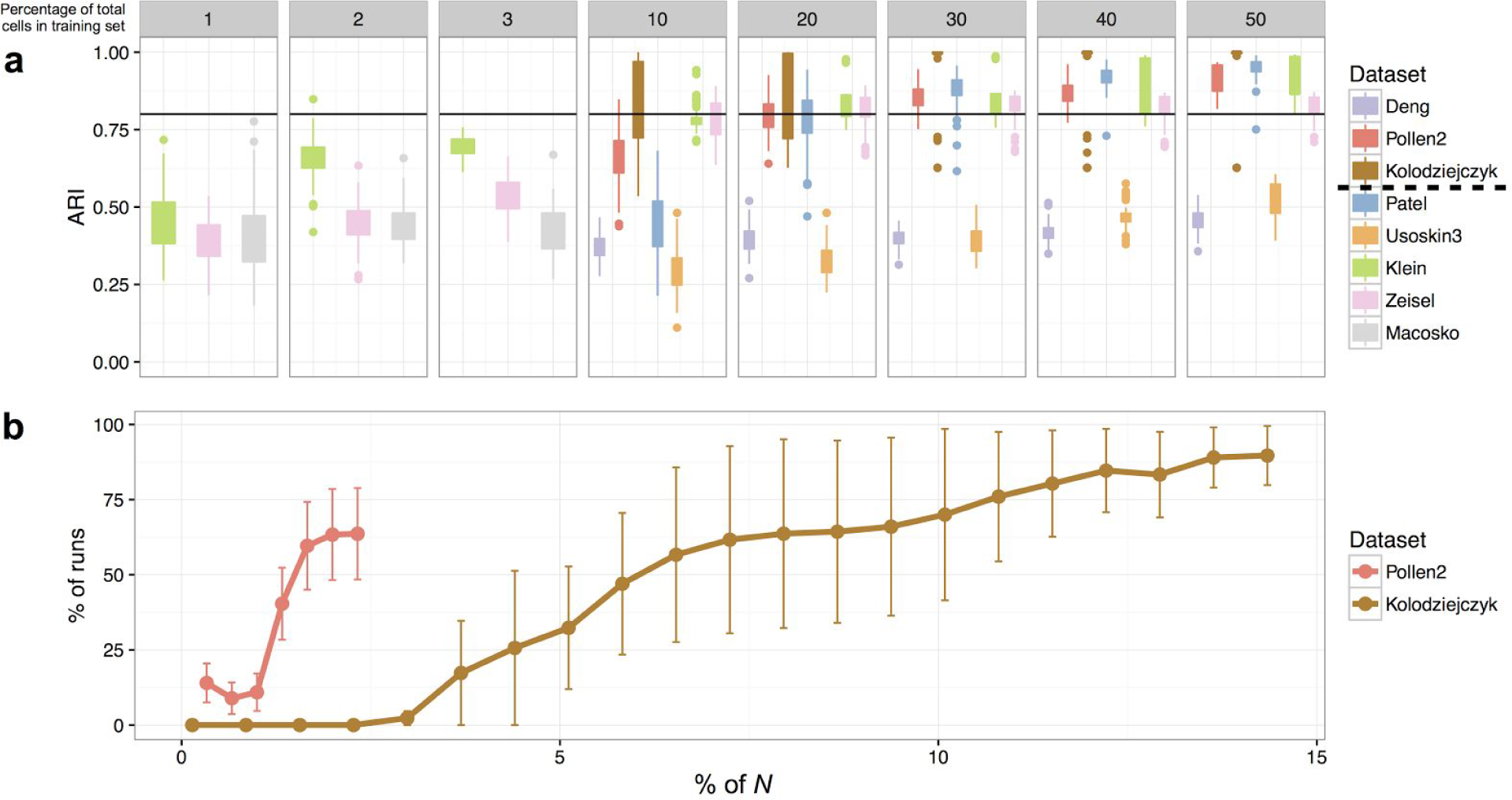
SC3 can be scaled to large datasets using a hybrid approach. **(a)** The performance of the hybrid SC3, as measured by the ARI, improves as the % of subsampled cells increases. The results indicate that accurate clustering can be achieved with only a small percentage of all cells used to obtain SC3 labels, which are then used as inputs by a linear kernel support vector machine (SVM). Dots represent outliers higher (lower) than the highest (lowest) value within 1.5 × IQR, where IQR is the interquartile range. The black line indicates ARI = 0.8. The dashed black line in the legend separates gold and silver standard datasets. (**b**) Robustness of SC3 for the detection of rare celltypes. For two of the datasets, we remove different percentages of the cells in the rare celltypes. The figure shows the mean fraction of SC3 runs in which all the rare cells were clustered together as a function of the total number of cells in the rare celltype.

To specifically evaluate the sensitivity of SC3 for identifying rare cell-types, we carried out a synthetic experiment, whereby cells from one cell-type were removed iteratively from the Kolodziejczyk and Pollen datasets (Methods). For the Pollen dataset, SC3 can detect clusters containing ˜1% of the cells, whereas for the Kolodziejczyk dataset ˜10% of the cells are required (Fig. 3b). Note that the ARI is not a useful metric here since the penalty for misclassifying only a few cells is very small, so instead we ask whether all of the rare cells are identified together in a separate cluster (Methods). We hypothesize that the ability to identify rare cells reflects the origins of the two datasets; the Pollen data is more diverse as it represents 11 different cell lines while the Kolodziejczyk data comes from one cell-type grown in three different conditions.

The main drawback of the sampling strategy is that we may fail to identify rare cell-types, and when *N*>>5,000 there is a substantial risk that the sampled distribution will differ significantly from the full distribution. When the hybrid SC3 approach is used, it is possible to exactly calculate the probability of missing out on a cell-type using the hypergeometric distribution if the proportion of sampled cells and the number of rare cells is known (Methods and Fig. S12). If the user is trying to identify a rare subpopulation (e.g. cancer stem cells), then the hybrid strategy is not recommended, and methods specifically designed to identify rare cell-types such as RaceID^1^ or GiniClust^26^ may be more appropriate.

### SC3 features an interactive and user-friendly interface

SC3 is implemented in R and as it is part of BioConductor^27^, it is easy to download, install and integrate into existing bioinformatics pipelines. To increase its usability, SC3 features a graphical user interface displayed through a web browser, thus minimizing the need for bioinformatics expertise. The user is only required to provide the input expression matrix and a range for the number of clusters, *k*. SC3 will then calculate the possible clusterings for this range. To help the user identify a good choice of *k*, we have implemented a method based on Random Matrix Theory (RMT)^28,^29^^ for determining the number of clusters. Briefly, the number of clusters is given by the number of eigenvalues of the normalized covariance matrix that differ significantly from the Tracy-Widom distribution (Methods). Overall, we find good agreement between these estimates, 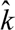, and the numbers suggested by the original authors (Fig. 2b). Furthermore, to aid the selection of a good clustering, SC3 also calculates the silhouette index^30^, a measure of how tightly grouped the cells in the clusters are. Since the clustering is performed during startup, the user can explore different choices of *k* in real time. The outcome is presented graphically as a consensus matrix to facilitate visual assessment of the clustering quality (Fig. 4a). The elements of the consensus matrix represent a score, normalised between 0 and 1, indicating the fraction of SC3 parameter combinations that assigned the two cells to the same cluster. By considering the consensus scores within and between clusters, one may quickly assess the clustering quality by visual inspection.

**Figure 4.**
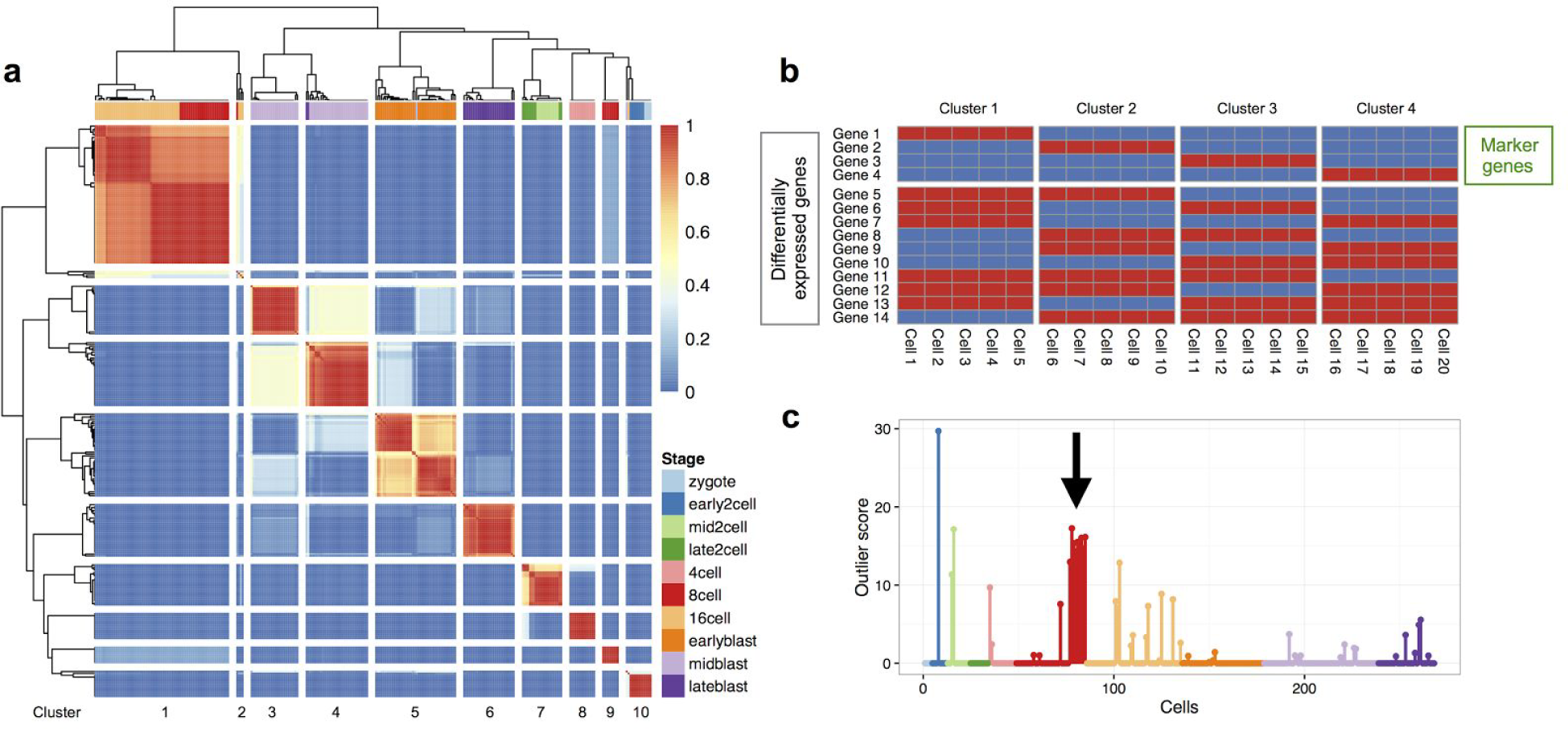
Applying SC3 to the Deng dataset aids biological interpretation. **(a)** The consensus matrix panel as generated by SC3. The matrix indicates how often each pair of cells was assigned to the same cluster by the different parameter combinations as indicated by the colorbar. Dark red (1) indicates that the cells were always assigned to the same cluster whereas dark blue **(0)** indicates that they were never assigned to the same cluster. In this case, SC3 finds a clustering with *k* = 10 clusters, separated by the white lines as visual guides. The colors at the top represent the reference labels, corresponding to different stages of development (see colour guide), **(b)** Illustration of the difference between marker genes and differentially expressed genes. In this small example, 20 cells containing **14** genes with binary expression values (blue for ‘off’, red for ‘on’) are clustered. Only genes **1-4** can be considered as marker genes, whereas all **14** genes are differentially expressed, **(c)** Outlier scores for all *N*= 268 cells as generated by SC3 (colors correspond to the 10 reference clusters provided by the authors - Stage in **(a)).** The nine cells with high outlier score in the red cluster (black arrow) were prepared using a different protocol (see text for details), and are thus assigned to a technical artifact.

### SC3 assists with biological interpretation

A key aspect to evaluate the quality of the clustering, which cannot be captured by traditional mathematical consistency criteria, is the biological interpretation of the clusters. To help the user characterise the biology of the clusters, SC3 identifies differentially expressed genes, marker genes, and outlier cells. By definition, differentially expressed genes vary between two or more clusters. To detect such genes, SC3 employs the Kruskal-Wallis^31^ test (Methods) and reports the differentially expressed genes across all clusters, sorted by *p*-value. The Kruskal-Wallis test has the advantage of being non-parametric, but as a consequence, it is not well suited for situations where many genes have the same expression value. Marker genes are highly expressed in only one of the clusters and are selected based on their ability to distinguish one cluster from all the remaining ones (Fig. 4b). To select marker genes, SC3 uses a binary classifier based on the gene expression to distinguish one cluster from all the others^15,32^. For each gene, a receiver operator characteristic (ROC) curve is calculated and the area under the ROC curve is used to rank the genes. The area under the ROC curve provides a quantitative measure of how well the gene can distinguish one cluster from the rest. The most significant candidates are then reported as marker genes (Methods). Cell outliers are also identified by SC3 through the calculation of a score for each cell using the Minimum Covariance Determinant33. Cells that fit well into their clusters receive an outlier score of 0, whereas high values indicate that the cell should be considered an outlier (Fig. 4c). The outlier score helps to identify cells that could correspond to, e.g., rare cell-types or technical artefacts. In addition, SC3 facilitates obtaining a gene ontology and pathway enrichment analysis for each cluster by directly exporting the list of marker genes to g: Profiler^34^. All results from SC3 can be saved to text files for further downstream analyses.

### SC3 can provide novel insights for published datasets

To illustrate the above features, we analysed the Deng dataset tracing embryonic developmental stages, including zygote, 2-cell, 4-cell, 8-cell, 16-cell and blastomere. Based on the silhouette index, a clustering into either *k*=2 groups or *k*=10 appears favorable, while the Random Matrix Theory based method recommends *k*=9 (Fig. 2b). The solution for *k*=2 identifies one cluster with the blastocyst and one with the remaining cells. The most stable result for *k*=10 is shown in Fig. 4a, and our clusters largely agree with the known sampling timepoints. However, our results suggest that the difference between the 8-cell and 16-cell stages is quite small. The latter stages of development are labelled “early”, “mid” and “late” blastomeres, although it is well known that these stages consist primarily of trophoblasts and inner cell mass. Interestingly, SC3 suggests that the mid-blastocyst stage could be split into two groups, which most likely correspond to trophoblasts and the inner cell mass (clusters 3 and 4). This conclusion is supported by the fact that *Sox2* and *Tdgf1* (inner cell mass markers) are listed as marker genes in cluster 4^35^ (Table S2). In total, we identified ˜3000 marker genes (Table S2), many of which had been previously reported as specific to the different developmental stages^36–40^. Furthermore, the analysis reveals several genes specific to each developmental stage which had previously not been reported (Table S2). Importantly, when using the reference labels reported by the authors^23^, nine cells have high outlier scores (red cells in Fig. 4c). As it turns out, these were prepared using the Smart-Seq2 protocol instead of the Smart-Seq protocol^12,23^, thus demonstrating the ability of our algorithm to identify outliers, which are introduced here ‘artificially’ due to the different technique used. Indeed, when we use SC3 to cluster the Deng data, the nine Smart-Seq2 cells form a separate cluster (#9 in Fig. 4a).

Using the hybrid approach, we are able to analyse a large Drop-Seq dataset with *N* = 44,808 cells and *k* = 39 clusters^15^ (Methods). The ARI between the SC3 clustering and the computationally-derived labels obtained by the original authors is 0.52. This result is largely driven by the fact that Macosko *et al*. lumped a large number of cells into a single “Rods” cluster. This Rods cluster contains 29,400 cells, but using SC3 a finer split of the Rods cluster is revealed with the majority of cells being assigned to 2 large clusters (clusters 4 and 8 on Fig 5). Interestingly, several genes related to photoreceptors (e.g. Gngt1, Pde6g, Rho, Rcvrn, Pdc, Gnat1, Nr1, Slc24a1, Rs1 and Sag for cluster 4; Rpgrip1 and Rp1 for cluster 8) are identified as markers distinguishing the two subclusters (Table S3), implying that there is likely a higher degree of heterogeneity amongst those cells than originally reported. We note that 94% of the 29,400 rod cells were lowly expressed (<900 genes detected), and this explains why so few marker genes were identified by SC3. Moreover, 31 of the clusters that were identified by SC3 can be matched with clusters identified by Macosko *et al*. (Fig 5 and Methods), suggesting that the subsampling employed in our hybrid strategy works well for larger datasets.

### SC3 characterises subclones in myeloproliferative neoplasm

Myeloproliferative neoplasms, a group of diseases characterised by the overproduction of terminally differentiated cells of the myeloid lineage, reflect an early stage of tumorigenesis where multiple subclones are known to coexist in the same patient^19^. From exome sequencing data, we previously identified TET2 and JAK2V61F as the only driver mutations in a large patient cohort^41^. In this paper, we use two patients out of this cohort which harbour TET2 and JAK2V617F mutations in their stem cell compartment. Haematopoietic stem cells (HSCs) are thought to be the cell of origin in myeloproliferative neoplasms. However, little is known about the transcriptional consequences of driver mutations on the stem cells. To gain more insight into the transcriptional landscape of patient derived HSCs, we obtained scRNA-seq data from the two patients (Methods). For patient 1(*N* = 51), both the silhouette index of SC3 and our RMT method suggested that *k* = 3, provides the best clustering, revealing three clusters of similar size (Fig. S17). For patient 2 (*N* = 89) SC3 indicated *k*=1, in agreement with the RMT algorithm, suggesting that one single cluster might best reflect the underlying transcriptional changes.

Since known driver mutations in these patients are the *TET2* and *JAK2V617F* loci^42^ we hypothesized that the different clusters correspond to different combinations of mutations within different clones. Unfortunately, the coverage of the *JAK2V617F* and *TET2* loci was insufficient to reliably determine the genotype of each cell directly from the scRNA-seq data. Instead the genotype composition for each HSC clone was determined by growing individual haematopoietic stem cells into granulocyte/macrophage colonies, followed by Sanger sequencing of the TET2 and JAK2V617F loci (Fig. 6a). In agreement with the clustering defined by SC3, patient 1 (*k*=3) was found to harbor three different subclones: (i) cells with both TET2 and JAK2V617F mutations, (ii) cells with a TET2 mutation and (iii) wild-type cells (Fig. 6b). Strikingly, the SC3-clusters contain 22%, 29% and 49% of the cells, in excellent agreement with the proportions of each genotype found in the patient, namely 20%, 30% and 50%. By contrast, none of the other clustering methods was able to clearly identify clusters which could readily be related to the genotype information (Supplementary Notes 3 and 4). Thus, we hypothesize that cluster 1 corresponds to the double mutant, cluster 2 corresponds to cells with only a TET2 mutation, and cluster 3 corresponds to wild-type cells. The HSC compartment of patient 2 was 100% mutant for TET2 and JAK2V617F, which again was consistent with clustering of *k*=1 suggested by SC3 (Fig. S18).

Four additional lines of evidence support the assumption that SC3 can help to define subclonal composition. Firstly, we analysed data from patient 2, with a dominant double mutant clone, together with all cells from patient 1, harbouring three different clones. SC3 clustering again suggested *k* = 3 (Fig. 5b, S19), in agreement with the RMT algorithm. Most importantly, all of the putative double mutant cells from patient 1 were grouped with the double mutant cells from patient 2. SC3 reported 33 marker genes for the putative *TET2* mutant and 202 marker genes for the putative double mutant clone (Fig. 5c, Table S5).

**Figure 5.**
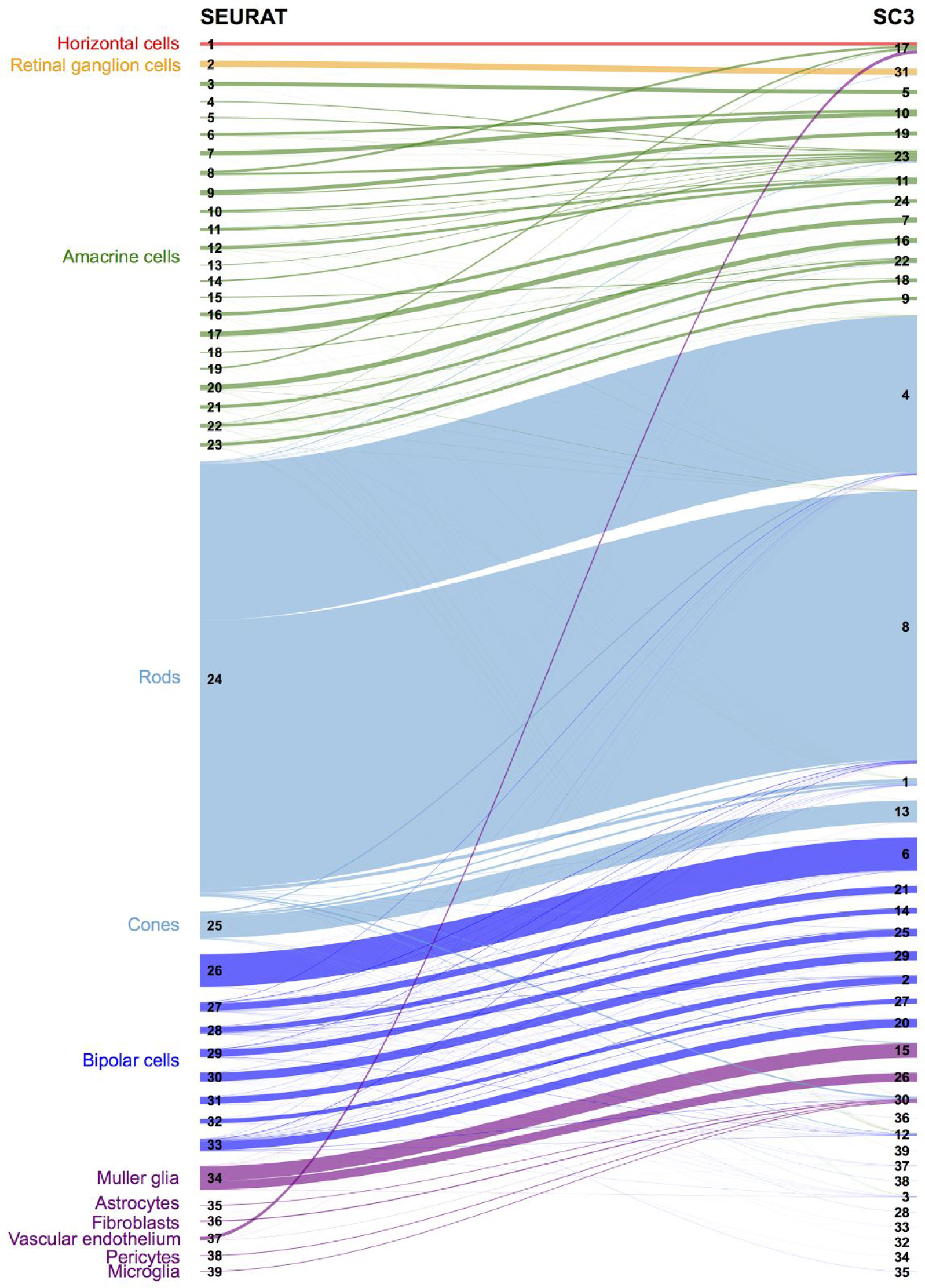
Analysis of SC3 clustering of the Macosko dataset. Sankey diagram comparing the 39 clusters reported by Macosko *et al*^15 15^ (left) and the 39 clusters obtained with SC3 (right). The widths of the lines linking both sets of clusters correspond to the number of cells they have in common. Colors and cell types as in ^15^.

**Figure 6.**
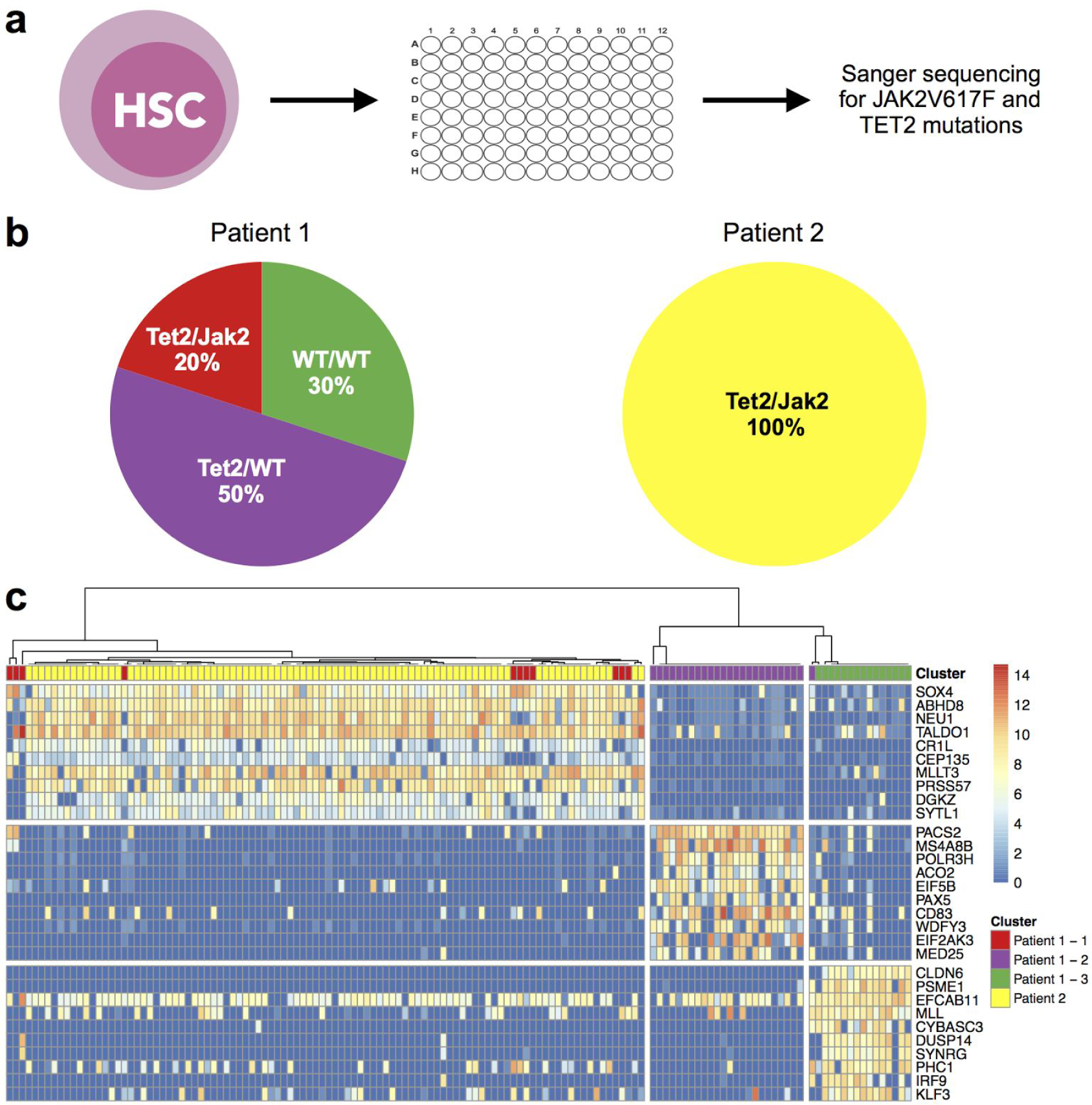
Using SC3 to define subclones from two patients with myeloproliferative neoplasm. **(a)** Individual HSCs were placed into wells, grown into granulocyte/macrophage colonies, and the *JAK2V617F* and the *TET2* loci were characterised using Sanger sequencing, **(b)** Clonal composition of patients 1, 2 obtained by independent sequencing experiments as described in Fig 6a of the *JAK2V617F* and the *TET2* loci (Methods), **(c)** Marker gene expression (after Gene Filter and Log-transformation, Methods) of the combined dataset (patient 1 + patient 2). Clusters (separated by white vertical lines) correspond to *k* = 3 (Methods). Cells corresponding to patient 1 are indicated with the same colour as in panel **(b).** Cells from patient 2 are indicated in yellow. Only the top 10 marker genes are shown for each cluster.

Secondly, we used microarray data from erythroid burst-forming units colonies available for patient 1^41^ where the genotype of each clone was linked to a specific transcriptional signature. When comparing differentially expressed genes for the double mutant clone from erythroid burst-forming unit colonies and the marker genes obtained from the pooled putative TET2/JAK2 mutant clone, we found 13 genes in common. This overlap was significant (*p*-value=0.048, hypergeometric test) and we also found a weak correlation (Spearman’s rho = 0.15, *p*-value=0.031) between the fold changes from the microarray and the scRNA-seq data.

Thirdly, we performed Gene and Pathway Enrichment Analysis using the marker genes (Methods). We found several categories related to haematopoiesis (selected with green color in Table S5). Among the enriched pathways were ‘Jak-STAT signalling pathway’, ‘estrogen signalling pathway’^43^ and ‘GPVI-mediated activation cascade’ (the latter plays a role in activation and aggregation of platelets). Furthermore, Gene Ontology analysis showed enrichment for the ‘Regulation of cytokine production’ term. Cytokines play an important role in haematopoiesis by initiating intracellular signals that govern cell fate choices such as proliferation and differentiation^19^. This confirms that ligands and receptors involved in JAK/STAT pathway activation are highly enriched in our marker genes for the putative double mutant cluster. For the putative TET2 only mutant subclones, none of the above pathways were specifically misregulated. Instead, we hypothesized that since *TET2* is involved in DNA de-methylation there would be a global impact on the transcriptome. Loss of TET enzymes has been reported to impact on the variability in gene expression in mouse embryos^37^. Comparing the genome-wide distribution of the normalized variances revealed that the putative TET2 mutants have more variable transcriptomes than putative wild-type cells (Mann-Whitney test *p*-value <2.2e-16, Methods and Fig. S20).

Fourth, SC3 identified several surface receptors from the list of marker genes corresponding to the different putative clusters. In particular, CD82 (corresponding to the putative double mutant), CD83 (WT clone) and CD127 or CD244 (Tet2 mutant clone) are surface markers that can be targeted by readily available, well-characterized commercial antibodies. We therefore carried out cell-sorting using such antibodies, and as predicted, only CD82 antigen, predicted to isolate cells with a double mutant nature, was expressed on the surface of CD34^+^CD38^+^cells from patient 2 (Fig. S21). In contrast, CD127 and CD83 antibodies were unable to isolate distinct cell populations from the same patient, strengthening the assumption that SC3 can predict clonal composition by providing specific marker genes. Due to limited material availability, we were only able to test one surface marker for patient 1. We chose CD244 since it was highly expressed in the putative Tet2 only mutant clone. (Fig. S21). Again, we were able to isolate a CD244 positive population in a subset of CD34^+^CD38^+^cells. This result demonstrates that SC3 is capable of characterising clusters defined by mutations rather than by patient batch.

## Discussion

We have presented SC3, an interactive tool for unsupervised clustering of scRNA-seq data. The central contribution of our work is the identification of a highly favorable regime for the number of dimensions to use for the clustering and the observation that a consensus strategy can improve both the stability and the accuracy of the clustering at a moderate increase of the computational cost. Interestingly, the favourable regime appears to be insensitive to reduced sequencing depth, and it is independent of how the data was normalized since it is similar for FPKM, TPM and UMI counts. It is likely that for samples with a different technical noise profile, or from a different genome size, this range might no longer be optimal. We hypothesize that the optimal range of *d* is related to the number of clusters, *k*. Assuming that the distance matrix can be written as ***A* =*X*** + ***Y***, where ***X*** is a random matrix of rank *N*, and ***Y***, representing the underlying biological signal, has rank *r<<N*, then we expect a gap in the eigenvalue spectrum of **A** after the *r*^th^ largest eigenvalue^44^. Consequently, we expect that using more than *r* eigenvectors will be suboptimal as the additional dimensions will represent the noise in ***X***. Similarly, if too few eigenvectors are used, then we will be unable to capture the structure of ***Y***. Thus, a sensible rule of thumb is to choose an interval of *d* values close to the ratio of 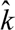/*N*. Interestingly, except for the Kolodziejczyk dataset, the 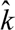/*N* ratio is in the range 3.7-7.8% of *N* for the gold standard datasets.

By comparing to several other methods, we demonstrate that SC3 provides a highly accurate clustering for published datasets. For large datasets, SC3 employs a hybrid approach, which makes it possible to scale the method to very large experiments, e.g. Drop-seq^15^. Importantly, SC3 features a graphical user interface, making it interactive and user-friendly. SC3 also aids biological interpretation by identifying outlier cells, differentially expressed genes and marker genes. Identification of differentially expressed genes for scRNA-seq data is currently an active area of research^32,45,46^, but these methods all require pairwise comparisons. Since the number of comparisons required is equal to *k(k-1)/2*, these methods are computationally costly to apply. Thus, developing methods for comparing more than two groups of cells is an important future direction of research.

A major challenge when developing unsupervised clustering algorithms for scRNA-seq is the lack of good mathematical models that can be used to generate realistic, synthetic surrogate datasets to benchmark the methods. Instead, we must rely on published datasets where the labels have been provided by the original authors. For some of the datasets (Biase, Yam, Goolam, Deng, Pollen and Kolodziejczyk), the labels are likely to be accurate since they correspond to cells taken from different tissues, conditions or time-points. For the other datasets, however, the labelling was based on a combination of the authors’ clustering methods and their biological knowledge. In these latter cases, the labelling is less reliable, and we cannot be certain that the original clustering represents a meaningful ground truth.

Although significant progress has been made on understanding the mutations leading to cancer^19^, much less is known about the differences between subclones within the tumour at the transcriptional level. We applied SC3 to scRNA-seq data from two patients diagnosed with myeloproliferative neoplasms. We found strong evidence in support of the hypothesis that the clusters revealed by SC3 directly correspond to subclones identified by independent experiments^41^. Moreover, we used the marker genes identified by SC3 to provide a biological characterisation of the different subclones (Fig. 6b, c). Amongst the marker genes reported by SC3 were several cell surface receptors which we were able to test using flow cytometry (Fig. S21). Our results demonstrate that it is possible to identify subclones using scRNA-seq with a high degree of confidence, and that the analysis of the transcriptome can provide important insights regarding the functional consequences of different mutations.

As sequencing costs decrease, larger scRNA-seq datasets will become increasingly common, furthering their potential to advance our understanding of biology. An exciting aspect of scRNA-seq is the possibility to address fundamental questions that were previously inaccessible, e.g. *de novo* identification of cell-types. However, the current lack of computational methods for analysing scRNA-seq has made it difficult to exploit fully the information contained in such datasets. We have shown that SC3 is a versatile, accurate and user-friendly tool, which will facilitate the analysis of complex scRNA-seq datasets. We believe that SC3 can provide experimentalists with a hands-on tool that will help extract novel biological insights from such rich datasets.

## Methods

### Gold standard datasets

All gold standard datasets (Fig. 1c), except the Pollen dataset, were acquired from the accessions provided in the original publications. The Pollen dataset^17^ was acquired from personal communication with Alex A Pollen.

### SC3 clustering

SC3 takes as input an expression matrix *M* where columns correspond to cells and rows correspond to genes/transcripts. Each element of M corresponds to the expression of a gene/transcript in a given cell. By default SC3 does not carry out any form of normalization or correction for batch effects. SC3 is based on five elementary steps. The parameters in each of these steps can be easily adjusted by the user, but are set to sensible default values, determined via the Biase, Deng, Yan, Goolam, Pollen and Kolodziejczyk datasets.

#### 1. Gene filter

The gene filter removes genes/transcripts that are either expressed (expression value is more than 2) in less than X% of cells (rare genes/transcripts) or expressed (expression value is more than 0) in at least (100-X)% of cells (ubiquitous genes/transcripts). By default X is 6. The motivation for the gene filter is that ubiquitous and rare genes are most often not informative for the clustering. We also explored all three parameters defined in the gene filter (expression thresholds of rare and ubiquitous genes/transcripts and the percentage X) and found that in general the gene filter did not affect the accuracy of clustering (Fig. S7). However, the gene filter significantly reduced the dimensionality of the data, thereby speeding up the method.

For further analysis the filtered expression matrix ***M*** is log-transformed after adding a pseudo-count of 1: ***M’*** = log2(***M*** + 1).

#### 2. Distance calculations

Distance between the cells, i.e. columns, in ***M**’* are calculated using the Euclidean, Pearson and Spearman metrics to construct distance matrices.

We investigated the impact of dropouts on distance calculations by considering a modified distance metric that ignores dropouts. This was done by excluding genes that were not expressed in at least one cell from the distance calculation. We found that this did not improve the performance (Fig. S9).

#### 3. Transformations

All distance matrices are then transformed using either principal component analysis (PCA) or by calculating the eigenvectors of the associated graph Laplacian (***L* = *I* - D**^−½^**AD**^−½^, where ***I*** is the identity matrix, ***A*** is a similarity matrix (***A*** = exp(−**A’**/max(**A’**))), where ***A’*** is a distance matrix) and ***D*** is the degree matrix of ***A***, a diagonal matrix which contains the row-sums of ***A*** on the diagonal (***D_ii_*** = **Σ*_j_A_ij_***). The columns of the resulting matrices are then sorted in descending order by their corresponding eigenvalues.

#### 4. k-means

*k*-means clustering is performed on the first *d* eigenvectors of the transformed distance matrices (Fig. 1a) by using the default kmeans() R function with the Hartigan and Wong algorithm^47^. By default, the maximum number of iterations is set to 10^9^ and the number of starts is set to 1,000.

#### 5. Consensus clustering

SC3 computes a consensus matrix using the Cluster-based Similarity Partitioning Algorithm (CSPA)^48^. For each individual clustering result a binary similarity matrix is constructed from the corresponding cell labels: if two cells belong to the same cluster, their similarity is 1, otherwise the similarity is 0 (Fig. 1a). A consensus matrix is calculated by averaging all similarity matrices of individual clusterings. To reduce computational time, if the length of the *d* range (*D* on Fig. 1a) is more than 15, a random subset of 15 values selected uniformly from the *d* range is used.

The resulting consensus matrix is clustered using hierarchical clustering with complete agglomeration and the clusters are inferred at the *k* level of hierarchy, where *k* is defined by a user (Fig. 1a). In principle, the *k* used for the hierarchical clustering need not be the same as the *k* used in step 5. However, for simplicity in SC3 the two parameters are constrained to have the same value.

Fig. 1e shows how the quality and the stability of clustering improves after *consensus clustering.*

### Adjusted Rand Index

If cell-labels are available (e.g. from a published dataset) the Adjusted Rand Index (ARI)^49^ can be used to calculate similarity between the SC3 clustering and the published clustering. ARI is defined as follows. Given a set of *n* elements, and two clusterings of these elements the overlap between the two clusterings can be summarised in a contingency table, where each entry denotes the number of objects in common between the two clusterings. The ARI can then be calculated as: 

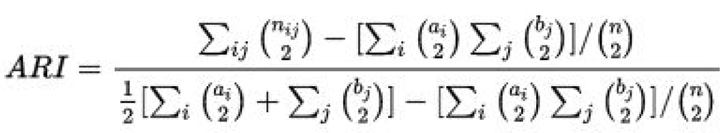

where *n_ij_* are values from the contingency table, *α_i_* is the sum of the *i*^th^ row of the contingency table, *b_j_* is the sum of the *j*^th^ column of the contingency table and () denotes a binomial coefficient.

Since the reference labels are known for all published datasets, ARI is used for all comparisons throughout the paper.

### Downsampling of the gold standard datasets

For each gene *i* and each cell *j*, the downsampled expression value was generated by drawing from a binomial distribution with parameters *p* = .1 and *n* = round(***M**_ij_*).

### Identification of a suitable number of groups 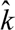

Matrix **Z** is obtained from *M*’ by subtracting the mean and dividing by the standard deviation for each column (z-score). Next, the eigenvalues of ***X = Z^T^*Z*** are calculated. The number of clusters 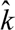 is determined by the number of eigenvalues that are significantly different with a p-value <.001 from the Tracy-Widom distribution^28,29^ with mean 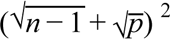 and standard deviation 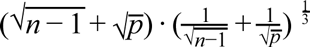, where *n* is the number of genes/transcripts and *p* is the number of cells.

### Benchmarking

For each dataset we used the expression units provided by the authors. The gene filter was applied to all the datasets. For tSNE+k-means, SNN-Cliq and pcaReduce the same log-transformation as in SC3 (***M***’ = log2(***M*** + 1)) was applied. For SINCERA we used the original z-score normalisation^16^ instead of the log-transformation. For tSNE the Rtsne R package was used with the default parameters. For SEURAT we used the original Seurat R package: we performed tSNE embedding with the default parameters once (following the authors’ tutorial at http://www.satijalab.org/clustertutorial1.html) and then clustered the data using DBSCAN algorithm multiple times, where we varied the density parameter *G* in the range 10^−3^−10^3^ to find a maximal ARI (this ARI is presented in Fig. 2). SEURAT was not able to find more than one cluster for the smallest datasets (Biase, Yan, Goolam, Treutlein and Ting) leading to very small ARI scores.

### Support Vector Machines (SVM)

When using SVM a specific fraction of the cells is selected at random with uniform probability. Next, a support vector machine^50^ model with a linear kernel is constructed based on the obtained clustering. We used the *svm* function of the *e1071* R-package with default parameters. The cluster IDs for the remaining cells are then predicted by the SVM model.

### Identification of rare cell-types

For the Pollen dataset, all but 1-7 of the cells in one of the 11 clusters were removed. The limit of 7 cells corresponds to the size of the smallest cluster in the original data. Subsequently, SC3 was run using *k*= 11, and we asked whether or not the cells of the rare cell-type were located in a separate cluster. This was repeated 100 times for each cell-type and Fig. 3b reports the percentage of runs when the rare cells were found together in a cluster with no other cells. Note that the ARI is a poor indicator of the ability to identify rare cells since this measure is relatively insensitive to the behavior of a small fraction of the cells. For the Kolodziejczyk dataset, we used a similar strategy, but we allowed for 1-101 cells in the rare group.

For the hybrid SC3 approach with 30% of cells used to train the SVM we were able to calculate the probability of including the rare cell-types in the training set analytically by multiplying the data from Fig. 3b by the probability of all rare cells to be included in the drawn sample (30% of all cells). This probability was calculated using the hypergeometric distribution R function: phyper(n.rare.cells -1, n.rare.cells, n.other.cells, 0.3*(n.other.cells + n.rare.cells), lower.tail=F), where n.rare.cells is the number of rare.cells and n.other.cells is the number of other cells in the dataset.

### Biological insights

#### Identification of differential expression

Differential expression is calculated using the non-parametric Kruskal-Wallis test^31^, an extension of the Mann-Whitney test for the scenario when there are more than two groups. A significant p-value indicates that gene expression in at least one cluster stochastically dominates one other cluster. SC3 provides a list of all differentially expressed genes with *p*-values <0.01, corrected for multiple testing (using the default “holm” method of p.adjust() R function) and plots gene expression profiles of the 50 most significant differentially expressed genes. Note that the calculation of differential expression after clustering can introduce a bias in the distribution of *p*-values, and thus we advise to use the *p*-values for ranking the genes only.

#### Identification of marker genes

For each gene a binary classifier is constructed based on the mean cluster expression values. The area under the receiver operating characteristic (ROC) curve is used to quantify the accuracy of the prediction. A *p*-value is assigned to each gene by using the Wilcoxon signed rank test comparing gene ranks in the cluster with the highest mean expression with all others (*p*-values are adjusted by using the default “holm” method of p.adjust() R function). The genes with the area under the ROC curve (AUROC)>0.85 and with the *p*-value<0.01 are defined as marker genes. The AUROC threshold corresponds to the 99% quantile of the AUROC distributions obtained from 100 random permutations of cluster labels for all datasets (Table S1 and Fig. S13). SC3 provides a visualization of the gene expression profiles for the top 10 marker genes of each obtained cluster.

#### Cell outlier detection

Outlier cells are detected by first taking an expression matrix of each individual cluster (all cells with the same labels) and reducing its dimensionality using the robust method for PCA (ROBPCA)^51^. This method outputs a matrix with *N* rows (number of cells in the cluster) and *P* columns (retained number of principal components after running ROBPCA). SC3 then uses p=min(*P*, 3) first principal components for further analysis. If ROBPCA fails to perform or *P*= 0, SC3 shows a warning message. We found (results not shown) that this usually happens when the distribution of gene expression in cells is too skewed towards 0. Second, robust distances (Mahalanobis) between the cells in each cluster are calculated from the reduced expression matrix using the minimum covariance determinant (MCD)^33^. We then used a threshold based on the *Q*% quantile of the chi-squared distribution (with *p* degrees of freedom) to define outliers. By default *Q*=99.99, but can be manually adjusted by a user. Finally, we define an outlier score as the difference between the square root of the robust distance and the square root of the *Q*% quantile of the chi-squared distribution (with *p* degrees of freedom). The outlier score is plotted as a barplot (Fig. 4c).

### Analysis of the Macosko dataset

To analyze the Drop-Seq dataset we followed the procedure used by Macosko et al^15^ and selected the 11,040 cells where more than 900 genes were expressed. Moreover, due to the low read depth, the gene filter was removed. We then sampled 5,000 cells and clustered using SC3, including the SVM step, 100 times. All 100 solutions were consistent between each other resulting in an average ARI of 0.6 and they were sufficiently accurate compared to the reference authors’ clustering yielding an average ARI of 0.54 (Fig. S14). Since each of the 100 solutions were different, we added an additional consensus clustering step using the “best of k” consensus algorithm^52^. This approach provided a single solution based on the 100 different solutions and it was as accurate as the individual solutions with an ARI of 0.52 (the actual labels are presented in Table S3). The SC3 consensus solution splits the large original cluster (cluster 24 with 29,400 cells) hierarchically into 2 clusters of smaller sizes (18105 + 10558 = 28663 cells, clusters 4 and 8 in Fig. 5). Additional gene and pathway enrichment analysis for the differentially expressed genes between the two clusters is presented in the Supplementary Note 5. If more than 75% of the cells from the reference cluster are shared with the SC3 cluster we defined these two clusters as matched. In total 31 reference clusters were matched to the SC3 clusters.

### Patients

Both patients provided written informed consent. Diagnoses were made in accordance with the guidelines of the British Committee for Standards in Haematology.

#### Isolation of haematopoietic stem and progenitor cells

Cell populations were derived from peripheral blood enriched for haematopoietic stem and progenitor cells (CD34+, CD38−, CD45RA−, CD90+), hereafter referred to as HSCs. For single cell cultures, individual HSCs were sorted into 96-well plates (Fig. S15) and grown in a cytokine cocktail designed to promote progenitor expansion as previously described^53^. For scRNA-seq studies, single HSCs were directly sorted into lysis buffer as described in Picelli et al^54^.

#### Determination of mutation load

Colonies of granulocyte/macrophage composition were picked and DNA isolated for Sanger sequencing for JAK2V617F and TET2 mutations as previously described by Ortmann *et al*^41^.

#### Single cell RNA-Sequencing

Single HSCs were sorted into 96-well plates and cDNA generated as described previously^54^. The Nextera XT library making kit was used for library generation as described by Picelli *et al*^54^.

#### Processing of scRNA-seq data from HSCs

96 single cell samples per patient with 2 sequencing lanes per sample were sequenced yielding a variable number of reads (*mean* = 2,180,357, *std dev* = 1,342,541). FastQC^55^ was used to assess the sequence quality. Foreign sequences from the Nextera Transposase agent were discovered and subsequently removed with Trimmomatic^56^ using the parameters HEADCROP:19 ILLUMINACLIP:NexteraPE-PE.fa:2:30:10 TRAILING:28 CROP:90 MINLEN:60 to trim the reads to 90 bases before being mapped with TopFlat^57^ to the Ensembl^58^ reference genome version GRCh38.77 augmented with the spike-in controls downloaded from the ERCC consortium^59^. Counts of uniquely mapped reads in each protein coding gene and each ERCC spike-in were calculated using SeqMonk (http://www.bioinformatics.bbsrc.ac.uk/projects/seqmonk) and were used for further downstream analysis. Quality control of the cells contained two steps: 1. filtering of cells based on the number of expressed genes; 2. filtering of cells based on the ratio of the total number of ERCC spike-in reads to the total number of reads in protein coding genes. Filtering threshold were manually chosen by visual exploration of the quality control features (Fig. S16). After filtering, 51 and 89 cells were retained from patient 1 and patient 2, correspondingly. The expression values in each dataset were then normalised by first using a size-factor normalisation (from DESeq2 package^60^) to account for sequencing depth variability. Secondly, to account for technical variability, a normalisation based on ERCC spike-ins was performed using the RUVSeq package^61^ (RUVg() function with parameter k = 1). For combined patient data, normalisation steps were performed after pooling the cells. The resulting filtered and normalised datasets were clustered by SC3. Potential biases of cell filtering on the proportions of cells in the clusters of patient 1 are considered in the Supplementary Note 1. It shows that the cluster of lower cell quality is separated from the other biologically meaningful clusters of patient 1 and it does not change the total proportion of the biologically meaningful clusters. Supplementary Note 2 shows that SC3 results of clustering of patient 1 do not depend on the normalization procedure.

#### Clustering of patient scRNA-seq data by SC3

We clustered scRNA-seq data from patient 1 and patient 2 separately as well as a combined dataset containing data from patient 1 + patient 2. For patient 1, in agreement with the RMT algorithm, the best clustering was achieved for *k* = 3 (Fig. S17). Data from patient 2 was homogeneous and SC3 was unable to identify more than one meaningful cluster (Fig. S18), again in agreement with the RMT algorithm. For the combined dataset for patient 1 + patient 2 the best values of the silhouette index were obtained when *k* was 2 or 3 (Fig. S19). In both cases all of the cells from cluster 1 in patient 1 were grouped with the cells from patient 2. For *k*= 3 clusters 1 and 3 of patient 1 were also resolved. The RMT algorithm also provided *k* = 3 for the merged patient 1 + patient 2 dataset.

#### Comparison of clustering of patient 1 scRNA-seq data

Results of the clustering of the patient 1 data by other methods and their comparison to SC3 is presented in the Supplementary Notes 3 and 4.

#### Identification of differentially expressed genes from microarray data

The microarray data of patient 1 (EE50) was obtained from Array Express accession number E-MTAB-3086^41^. One replicate (2B) was identified as an outlier and removed. The limma R package^62^ was used to identify 932 differentially expressed genes between WT and TET2/JAK2V617F double mutant using an adjusted (by false discovery rate) *p*-value threshold of 0.1.

#### Marker genes analysis for patients

For both patients, to increase the number of marker genes, the AUROC threshold was set to 0.7 instead of the default value of 0.85 and the 0.1 *p*-value threshold was chosen.

#### Pathway enrichment analysis

We utilized g:Profiler web tool^34^ to perform gene and pathway enrichment analysis in obtained set of marker genes. The results are presented in Table S5.

## Contributions

MH conceived the study; VYK, MH, MTS, MB, TA and AY contributed to the computational framework; KK and TC performed the experiments for the patient data; KNN helped with the analysis of embryonic mouse data; MB, WR, ARG and MH supervised the research; VYK and MH led the writing of the manuscript with input from the other authors.

## Accession Numbers

scRNA-seq data for patient 1 and 2 is available from GEO accession GSE79102.

## Software availability

SC3 is available as a R package at http://bioconductor.org/packaaes/SC3/.

## Acknowledgements

We would like to thank Borislav Vangelov, Jean-Charles Delvenne and Renaud Lambiotte for fruitful discussions and their help with computational methods. We would also like to thank David Flores Santa Cruz, Danai Dimitropolou and Jacob Grinfeld for technical assistance with experiments. We thank Ignacio Vasquez-Garcia, David Harmin, Michal Kosicki, Daniel Ramsköld and Meri Huch for helpful comments on the manuscript.

